# Coupled organoids reveal that signaling gradients drive traveling segmentation clock waves during human axial morphogenesis

**DOI:** 10.1101/2022.05.10.491359

**Authors:** Yusuf Ilker Yaman, Roya Huang, Sharad Ramanathan

## Abstract

Axial development of mammals is a dynamic process involving several coordinated morphogenetic events including axial elongation, somitogenesis, and neural tube formation. How different signals control the dynamics of human axial morphogenesis remains largely unknown. By inducing anteroposterior symmetry breaking of spatially coupled epithelial cysts derived from human pluripotent stem cells, we were able to generate hundreds of axially elongating organoids. Each organoid was composed of a neural tube flanked by presomitic mesoderm that was sequentially segmented into somites. Periodic activation of the somite differentiation gene MESP2 coincided in space and time with anteriorly traveling segmentation clock waves in the presomitic mesoderm of the organoids, recapitulating key aspects of somitogenesis. Through timed perturbations of organoids, we demonstrated that FGF and WNT signaling play distinct roles in axial elongation and somitogenesis, and that the segmentation clock waves are driven by FGF signaling gradients. By generating and perturbing organoids that robustly recapitulate the architecture and dynamics of multiple axial tissues in human embryos, this work offers a means to dissect complex mechanisms underlying human embryogenesis.

## Introduction

The progenitors in the tailbud of the axially elongating mammalian embryo give rise to the posterior neural tube and the flanking presomitic mesoderm (PSM) (Gouti et al., 2017; Hubaud and Pourquie, 2014). The PSM is further patterned and segmented into somites (Fig 1A), which in turn give rise to the axial skeleton, body skeletal muscles and dorsal dermis (Pourquie, 2011), while the posterior neural tube gives rise to the spinal cord (Gouti et al., 2015). Periodic and sequential segmentation of PSM into somites is controlled by the segmentation clock, which is a complex network of oscillating genes under the control of NOTCH, FGF and WNT pathways (Dequeant et al., 2006). The cyclic expression of these genes travels anteriorly through the PSM as a gene expression wave (Aulehla et al., 2003; Hayashi et al., 2009; Palmeirim et al., 1997). When each such segmentation clock wave reaches the anterior end of the PSM, it initiates the segmentation program of the next presumptive somite pair. Thus, the boundary between the somites and the undifferentiated PSM, called the somite determination front, moves posteriorly with every wave. During development, the embryo must coordinate the dynamics of multiple processes including axial elongation, PSM generation, anterior movement of the segmentation clock waves in the PSM, posterior movement of the somite determination front, and somite segmentation. FGF and WNT pathways have been shown to be required for axial elongation (Amin et al., 2016; Benazeraf et al., 2010; Dubrulle and Pourquie, 2004) and the differentiation of axial progenitors into presomitic mesoderm (Wymeersch et al., 2021) in mouse, chick, and zebrafish. In addition, the position of the somite determination front along the anteroposterior axis is thought to be defined by FGF and WNT signaling gradients in different vertebrates (Aulehla et al., 2008; Dubrulle et al., 2001; Dubrulle and Pourquie, 2004; Dunty et al., 2008; Naiche et al., 2011). However, the mechanisms underlying the anterior movement of segmentation clock waves and the interaction of these waves with the signaling gradients remain unknown (Diaz-Cuadros and Pourquie, 2021).

**Fig.1.**
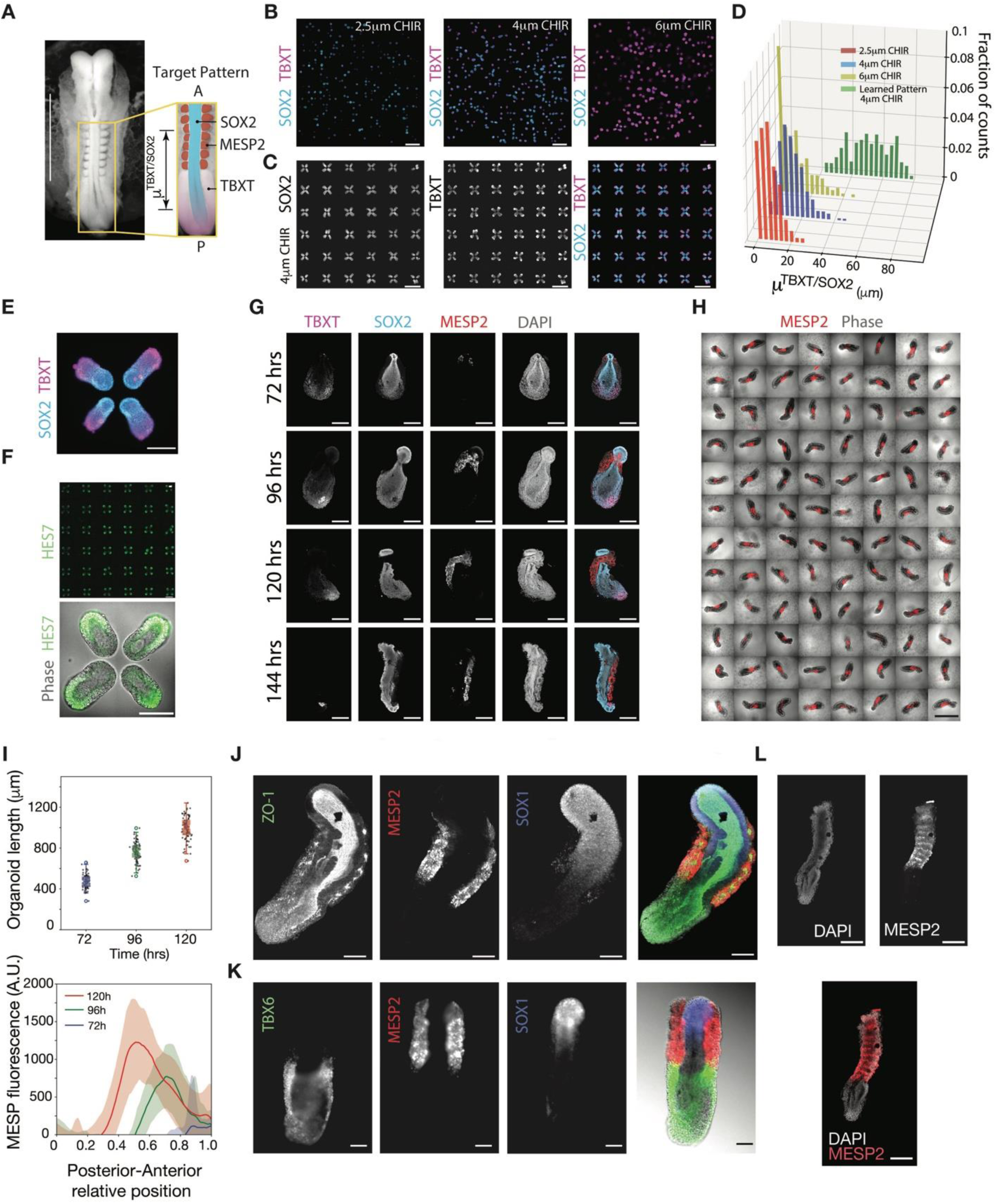
Elongating axial organoids generates neural tube with a single lumen flanked anteriorly by segmented somites and posteriorly by presomitic mesoderm. (**A**) Left: Micrograph of Carnegie Stage 10 human embryo, Kyoto Collection. Right: Target pattern with expression profiles of MESP2, TBXT and SOX2 colored and overlayed on posterior part of the human embryo micrograph. (**B**) Randomly positioned organoids on a coverslip, each consisting of a single epithelial layer of cells enclosing a single lumen, treated with BMP inhibitor LDN193189 (0.5µM), TGFβ inhibitor A83-10 (0.5µM) and WNT agonist CHIR99021 (left to right: 2.5µM, 4µM, 6µM) for 48 hours stained for TBXT and SOX2. Scale bar, 1mm. (**C**) Organoids micropatterned in groups of four on the vertices of a square after 48 hours of differentiation, stained for TBXT (left) and SOX2 (middle), color combined (right). Scale bars, 1 mm. (**D**) Histogram showing the distribution of organoid polarization metric on random arrays for each CHIR concentration and on the vertices of squares at 4µM CHIR concentration. (**E**) Magnified image of one set of four organoids on the vertices of a square at 48 hours of differentiation stained for TBXT and SOX2. Scale bar, 200µm. (**F**) Organoids micropatterned in groups of four on the vertices of a square after 48 hours of differentiation with live HES7 reporter signal (green) shown for a full array (top); magnified and overlayed on the phase image for one set of four organoids (bottom). One hundred percent of the organoids on coverslips show polarized expression HES7. Scale bars: 1mm top, 200µm bottom. (**G**) Confocal sections of representative organoids with MESP2::mCherry reporter on consecutive days of differentiation (72 h, 96 h, 120 h, 144 h) stained for TBXT, SOX2. TBXT and SOX2 co-expressing NMPs reside at the posterior tip. MESP2 progression starts at the anteriorly and moves towards the tip during 144 hours of differentiation. Scale bars, 200µm. (**H**) Phase contrast images overlayed with MESP2::mCherry signal in live organoids in a 96-well low adhesion plate at 120 h of differentiation. All organoids show elongated morphology and lateral MESP2::mCherry expression. Scale bar: 1mm. (**I**) Top: Quantification of elongation. Box plot of length of the organoids on 72h, 96h and 120h of differentiation. Dots are individual data points. Center line, median; box, interquartile range; whiskers, range not including outliers; empty circles: outliers. n= 95 independent biological replicates. Bottom: Quantification of MESP2 reporter expression along the anteroposterior axis of the organoids on 72h, 96h and 120h of differentiation. Solid lines: means; shaded area: std. AU, arbitrary units. n=95 independent biological replicates. (**J**) Confocal section of a representative organoid with MESP2:mCherry reporter stained for tight junction marker ZO-1 and SOX1 at 120h of differentiation. Scale bars: 100µm. (**K**) Epifluorescence image of a representative organoid with MESP2:mCherry reporter stained for paraxial mesoderm marker TBX6 and neural marker SOX1 at 120h of differentiation. Color combined fluorescence images overlayed on phase contrast image (right-most image). Scale bars: 100µm. (**L**) Confocal section of a representative organoid showing alternating expression of MESP2 reporter on 120 h of differentiation, stained for DAPI. Scale bar, 200µm.

The challenges in measuring and perturbing the dynamics of mouse embryos *in utero* and the ethical challenge in studying human development necessitate the use of *in vitro* systems. Recently, oscillating gene expression patterns and the generation of somitic mesoderm have been reported in monolayer cultures of human pluripotent stem cell-derived PSM cells, which is the first evidence for the existence of a segmentation clock in humans (Chu et al., 2019; Diaz-Cuadros et al., 2020; Matsuda et al., 2020). In exciting recent work, mouse and human stem cell-derived organoids have been directed to extend axially and generate neural progenitors and somites (Sanaki-Matsumiya et al., 2022; van den Brink et al., 2020; Veenvliet et al., 2020). However, the morphological variability from organoid to organoid and defects in the architecture of the underlying tissues in these *in vitro* systems significantly limit the use of chemical and genetic perturbations to gain mechanistic insight (Gupta et al., 2021). The ability to generate reproducible and robust organoids that recapitulate human axial development is therefore essential to make progress. Furthermore, such organoids will allow dynamic measurements and temporally controlled perturbations in a manner that is impossible *in vivo*.

Here, we aimed to characterize the dynamics of human axial patterning and morphogenesis to understand how segmentation clock waves interact with signaling gradients. Following the methods developed in a companion manuscript (Anand et al., 2022), we employed a combination of machine learning and bioengineering tools to tune the coupling of human pluripotent stem cell organoids by controlling their spatial arrangement. This enabled us to generate hundreds of *in vitro* organoids at a time, each of which robustly and reproducibly recapitulates the architecture of axial tissues in human embryos. Using single cell sequencing and computational analysis, we validated the organoids by determining the cell type composition as well as the spatial profiles of key transcription factors and signaling molecules along the anteroposterior axis. We demonstrated that the organoids recapitulate the dynamics of axial elongation, anteriorly moving segmentation clock waves, posteriorly moving somite determination front and somite segmentation. By perturbing the organoids, we showed that FGF signaling gradients drive the anterior propagation of segmentation clock waves, while simultaneously controlling the movement of the somite determination front, somite segmentation, and together with WNT, axial elongation. We finally discuss the implications of these results in the context of the existing models for axial patterning of the somites.

## Results

### Spatially coupled organoids achieve robust A-P symmetry breaking

To understand human axial development, we aimed to generate hundreds of organoids, each with an axially extending tailbud that generates a single lumen neural tube flanked posteriorly by PSM and anteriorly by somites as seen *in vivo* (Fig. 1A). To achieve this desired outcome, we built upon a bioengineering and machine learning framework developed in a companion manuscript, to reproducibly break anterior-posterior (A-P) symmetry (Anand et al., 2022). We first recapitulated the human epiblast (Zheng et al., 2019) by micropatterning human pluripotent stem cells at random locations on a glass coverslip and folding them into 150 µm diameter cysts composed of a single epithelial layer of pluripotent stem cells enclosing a lumen (Anand et al. 2022). To induce differentiation, we exposed the cysts to medium containing WNT agonist CHIR99021 (at concentrations ranging from 2.5µM to 6µM) while inhibiting BMP (LDN193189, 0.5µM) and TGF-beta signaling (A83-01, 0.5µM). After 48 hours, we stained the differentiated organoids for SOX2 and TBXT (Fig. 1B, Fig. S1A). These markers were chosen to label the SOX2+TBXT+ neuromesodermal (NMP) progenitors in the tailbud, SOX2-TBXT+ paraxial mesoderm flanking the neural tube posteriorly, and the SOX2+TBXT-cells of the neural tube. In the desired organoid morphology, we expected TBXT to be posteriorly expressed relative to SOX2. We thus scored our organoids using a polarization metric, 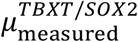, defined as the distance between the centroid of TBXT+ cells and that of SOX2+ cells in each organoid (Fig. 1A). The organoids on the random pattern showed a large variability in 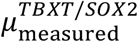 (Fig. 1B and D). Using the approach in Anand *et al*., we optimized the spatial arrangement of the differentiating organoids on a coverslip and the CHIR concentration such that each organoid broke A-P symmetry to acquire a large 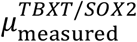. This arrangement consisted of 150µm diameter organoids in groups of four at the vertices of 200 by 200 µm squares (Fig. 1C, Fig. S1A). After exposure to 4µM CHIR for 48 hours, every organoid broke A-P symmetry and was polarized with low SOX2/ high TBXT posteriorly and high SOX2/ low TBXT anteriorly (Fig. 1C-E, Fig. S1A). To visualize this polarization in live organoids, we used a dual fluorescence reporter HES7:Achilles/MESP2:mCherry iPSC line (Diaz-Cuadros et al., 2020), allowing us to monitor the polarization of the presomitic mesodermal marker HES7 (Fig. 1F).

### Polarized organoids with posteriorly localized NMPs generate a single-lumen neural tube flanked by segmented somites

After the initial A-P symmetry breaking at 48 hours, organoids were removed from the coverslip and cultured individually in low adhesion 96 well plates in basal media (E6) supplemented with Matrigel, without any signaling molecules or inhibitors. From 72 to 120 hours after the start of differentiation, all the organoids that were successfully transferred to the 96 well plate underwent axial elongation (Fig. 1H and I, Fig. S1B). They also contained a TBXT+SOX2+ NMP population at the posterior tip that was maintained throughout 120 hours of differentiation (Fig. 1G, Fig. S1C). Every organoid showed lateral expression of the somitogenesis marker MESP2 based on mCherry expression (Fig. 1H) and displayed anterior to posterior progression of the MESP2+ somite fate (Fig 1H-I, Fig. S1B, n=95).

We next determined the expression patterns of key proteins in these organoids at 120h through immunostaining. Every organoid had an anteriorly positioned SOX1+ and SOX2+ neural tube with a single lumen and the proper apicobasal polarity as shown by ZO-1 and N-cadherin stains marking the apical tight junctions between epithelial cells (Fig 1J, Fig. S1F and G). In each of these organoids, the neural tube was flanked posteriorly by TBX6+ paraxial mesoderm, and anteriorly by the MESP2+ somite cells (Fig. 1K, Fig. S1D). ZO-1 and N-cadherin expression was localized in multiple foci in the MESP2+ somite region, showing the segmented architecture of somites flanking the anterior neural tube (Fig 1J, Fig S1F and G). Similar to the A-P organization of the developing embryo (Oginuma et al., 2008), the TBX6-expressing domain formed a clear boundary corresponding to the somite determination front, posterior to the MESP2+ cells (Fig. 1K, Fig. S1D and E). By 120 hours of differentiation, the MESP2 expression pattern had resolved into an alternating pattern in each segment as seen *in vivo* (Oginuma et al., 2008; Saga et al., 1997; Takahashi et al., 2000), indicating that the somites displayed A-P compartmentalization (Fig 1L). These results together demonstrate that our approach robustly achieved the differentiation of pluripotent stem cells into organoids with correctly positioned tailbud, neural tube, and somite structures, capturing key aspects of *in vivo* axial development.

### Clustering and diffusion map analysis of axial organoid transcriptome reveals cell types and A-P organization of neural tube and paraxial mesoderm

To explore the cell type composition of organoids, we performed single-cell RNA sequencing of 11009 cells obtained from 10 organoids at 120 h (Fig. S2A). Clustering and identifying cell types from this data requires a measure of distance between cells in gene expression space. The Euclidean distance in the space of all high variance genes leads to incorrect clustering and classification (Friedman et al., 2001). To overcome this challenge, we previously developed and validated an unsupervised statistical method, sparse multimodal decomposition (SMD), to identify the key subset of genes that can be used to determine cell types (Melton and Ramanathan, 2021). Using SMD, we identified 48 key genes with significant z-scores from the single cell data (Fig. 2A, Fig. S2B). Through hierarchical clustering in this gene subspace, we identified seven cell types in the organoids (Fig 2, A and B, Fig. S2, B and C). The first cluster co-expressed TBXT and SOX2, indicating a neuromesodermal progenitor identity (Gouti et al., 2017). The second cluster, consisting of SOX2+TBXT-cells, co-expressed tailbud genes HOXA10, CDX2, and NKX1-2, consistent with a pre-neural tube identity (Cooper et al., 2022). A third cluster possessed a neural progenitor identity, expressing neural markers SOX1, PAX6, HES5, and IRX3 along with high levels of SOX2 (Gouti et al., 2015). Three additional clusters were associated with paraxial mesodermal identity. These included a presomitic mesoderm-like cell cluster expressing TBX6, MSGN1, and HES7, early somite cell cluster expressing MEOX1, TCF15 and RIPPLY1, and a mature somite cell cluster expressing PAX3, TWIST1, and FST (Williams and Ordahl, 1994). Lastly, we identified a small number of notochord cells, expressing SHH, NOTO, and high levels of TBXT (Beckers et al., 2007; Resende et al., 2010).

**Fig.2.**
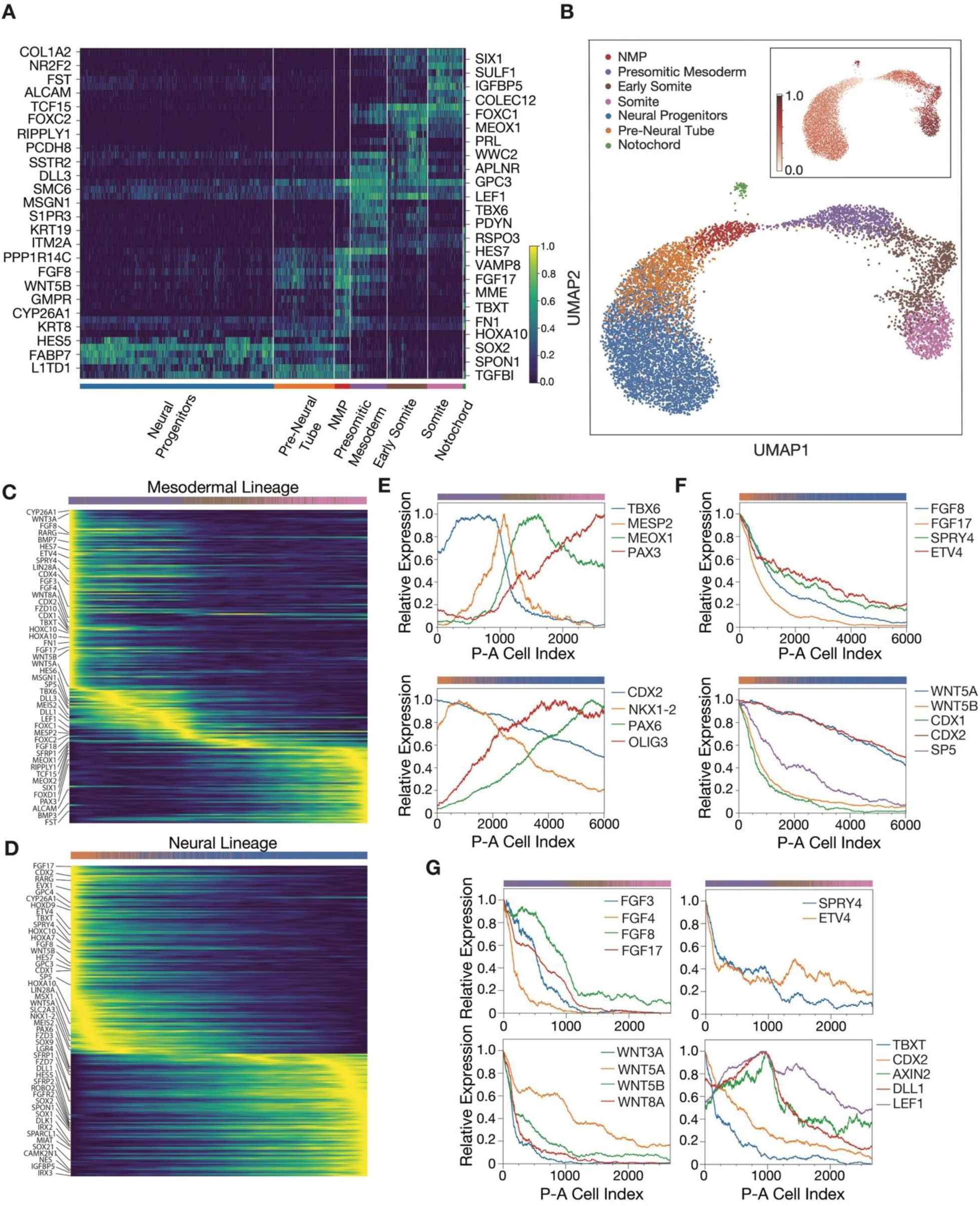
Anteroposterior organization of cell types and gene expression profiles inferred from single cell RNA-seq. (**A**) Normalized gene expression heatmap of 11009 cells from 120 h organoids, hierarchically clustered in the subspace of 48 genes identified by sparse multimodal decomposition. The inferred identities of the 7 clusters are labeled below. (**B**) UMAP (uniform manifold approximation and projection) plot generated in the subspace of genes identified by sparse multimodal decomposition. Cells are colored by their cluster identity (see legend for color code, same as in A). Inset shows the UMAP plot colored by inferred anteroposterior positions of cells obtained by diffusion mapping (see main text, methods). (**C** and **D**) Heatmap of top 200 differentially expressed genes (y axis) in the mesodermal (C) cell clusters (presomitic mesoderm, early somite, and somite) and neural (D) cell clusters (pre-neural tube and neural progenitors) in cells (x axis) ordered according to their inferred anteroposterior positions. Genes are ordered based on the position of their peak expression on the inferred A-P axis. Color bars on the top of heatmaps represent the cluster identity of the individual cells (same color code as in A). (**E, F**, and **G**) Normalized posterior-anterior gene expression profiles for marker genes of mesodermal (E, top) and neural cell clusters (E, bottom); FGF pathway ligands and targets (F, top), WNT pathway ligands and targets (F, bottom) in neural clusters; FGF pathway ligands (G, top left) and targets (G, top right), WNT pathway ligands (G, bottom left) and targets (G, bottom right) in mesodermal cell clusters. Bars on the top of each plot represents cells at that position along the inferred P-A axis, colored by their cluster identity (same color code as in A).

To map the spatial distribution of neural and mesodermal lineages present in organoids from scRNA-seq data, we constructed a diffusion map (Haghverdi et al., 2016) in the subspace of genes identified by SMD (Fig 2B). We previously showed that such a map can recapitulate the spatial ordering of cells along the A-P axis, allowing the inference of spatial profiles of gene expression (Anand et. al. 2022). We plotted the expression levels of genes arranged in order of the position of their peak expression level along the inferred posterior to anterior axis in both the neural and mesodermal tissues (Fig. 2, C and D). In the mesodermal cells, genes expressed in the tailbud progenitors (TBXT, CDX1, CDX2, CDX4, LIN28A, HOXA10, HOXC10) peak most posteriorly, followed by the marker genes of presomitic mesoderm (TBX6, DLL3, DLL1, MSGN1), followed by the somite determination front marker MESP2, then early somite markers (MEOX1, RIPPLY1, TCF15), and finally mature somite markers (SIX1, PAX3, ALCAM) (Grifone et al., 2005)(Fig. 2, C and E, Fig. S2D). Along the neural lineage, cells closest to the neuromesodermal progenitors expressed pre-neural tube markers CDX2, MSX1, and NKX1-2 (Fig. 2D, Fig. S2D). On the other hand, anteriorly positioned cells expressed higher levels of neural progenitor markers SOX1, PAX6, OLIG3 and IRX2, together with higher levels of SOX2 (Gouti et al., 2015)(Fig. 2, D and E, Fig. S2D). We verified the inferred anterior-posterior (A-P) expression profiles of the marker genes by MESP2:mCherry reporter expression and immuno-staining against TBXT, TBX6 (Fig. S1, C and D) and for mesodermal cells and SOX2, SOX1 and PAX6 for neural cells (Fig. S2H, S1C).

### Inferring the anteroposterior profiles of WNT, FGF, RA, NOTCH and BMP signaling pathway components in the mesodermal and neural tissues

We next tested whether our organoids recapitulated the anteroposterior expression gradients of WNT, FGF, RA, and NOTCH signals and their targets as observed *in vivo* in mouse, chick, and zebrafish embryos (Aulehla and Pourquie, 2010; Gouti et al., 2015). The expression profile of the detected FGF ligands in neural cells (FGF8, FGF17, Fig. 2F) and mesodermal cells (FGF3, FGF4, FGF8, FGF17, Fig. 2G) were localized most posteriorly in each lineage. FGF receptors were differentially expressed between the mesoderm and neural tissues, with FGFR1 expressed throughout the mesoderm and FGFR2 showing monotonically increasing levels from posterior to anterior in the neural tube (Fig. S2E). In both tissues, FGF target genes SPRY4 and ETV4 were upregulated posteriorly (Fig. 2, F and G), consistent with the role of FGF in maintaining the axial progenitor state in the tailbud of the mouse embryo (Diez del Corral et al., 2003; Naiche et al., 2011).

Similar to FGF, all detected WNT ligands were expressed in a posterior to anterior graded manner. While both canonical (WNT3A, WNT8A) and noncanonical (WNT5A, WNT5B) WNT ligands were expressed in the mesodermal tissue (Fig. 2G), only noncanonical (WNT5A, WNT5B) WNT ligands showed expression in the neural tissue (Fig. 2F). The WNT ligand expression gradient was opposed by an expression gradient of secreted WNT inhibitors SFRP1 and SFRP2 in both mesoderm and neural lineages (Fig. S2F). WNT targets (CDX2, CDX1, SP5) showed a posteriorly restricted expression pattern in the neural tissue similar to that of WNT ligands (Fig. 2F). In the mesodermal tissue, one class of WNT targets (TBXT, CDX2, CDX4), was highly expressed at the posterior end of the tissue and downregulated anteriorly. A second class of targets (AXIN2, DLL1) as well as the WNT transcriptional mediator LEF1, showed peak expression at the anterior end of presomitic mesoderm closest to the somite determination front (Fig. 2G).

Retinoic acid signaling is known to be important in mouse embryos for fate specification of NMPs, differentiation of presomitic mesoderm, and the patterning of the neural tube (Gouti et al., 2017). We observed that in our organoids, while the retinoic acid receptor gamma (RARG) was expressed throughout the neural and mesodermal tissues, the retinoic acid synthesis gene ALDH1A2 was expressed only in the somites anteriorly (Fig. S2G). Anterior retinoic acid secretion from somites combined with the posterior expression of the retinoic acid degradation enzyme CYP26A1 (Fig. S2G) is consistent with an A-P retinoic acid gradient. Anterior upregulation of transcription factor PAX6 known to be downstream of retinoic acid signaling (Sasai et al., 2014) suggested that the neural tube was patterned by retinoic acid secreted by the flanking somites. Consistent with this, immunostaining showed upregulation of PAX6 protein in the section of the neural tube in proximity to the somites (Fig. S2H).

Next, we investigated the NOTCH signaling pathway, a key component in the regulation of periodic somite segmentation and neurogenesis (Conlon et al., 1995; Moore and Alexandre, 2020). In the mesodermal lineage, both NOTCH pathway ligands (DLL1, DLL3), receptor (NOTCH1), and targets (HES6, HES7) were highly expressed in the presomitic mesoderm and downregulated anterior to the somite determination front (Fig. S2I). Although the NOTCH receptor expression levels were very low in the neural tissue, NOTCH ligand DLL1 and NOTCH target gene HES5 were upregulated in the neural progenitors anteriorly, suggesting the initiation of neurogenesis (Fig. S2I). In total, our human organoid model shows anteroposterior expression gradients of WNT, FGF, RA, and NOTCH signals and targets consistent with *in vivo* observations in mouse, chick, and zebrafish embryos.

Dorsoventral patterning of somites and neural tube is known to be regulated by opposing gradients of BMPs secreted by surface ectoderm, roof plate, and somites, and SHH secreted by the notochord and floor plate (Wilson and Maden, 2005). While our organoids lack roof plate and floor plate cell types, BMP7 was expressed by neuromesodermal progenitors and the posterior presomitic mesoderm, and BMP4 and BMP3 were expressed by somites (Fig. S2J). The presence of dorsalizing signals was consistent with the acquisition of a dorsal identity by both neural progenitor cells and somite cells. Neural progenitor cells expressed dorsal neural tube markers MSX1, OLIG3, IRX3 and PAX3 (Sagner and Briscoe, 2019), and somite cells expressed a dermomyotome marker PAX3 (Williams and Ordahl, 1994), seen only dorsally in somites *in vivo*. In contrast, ventral somite marker PAX1 (Ebensperger et al., 1995) was not expressed in any cells (Fig. S2J), consistent with the lack of SHH expression by cells. We also observed the expression of a neural crest marker SOX9, along with the epithelial-to-mesenchymal transition gene SNAI2 in a subset of neural progenitors (Fig. S2K). Thus, in the absence of SHH, tailbud progenitors generated only dorsal neural and mesodermal cell types, possibly through the dorsalizing effect of BMPs secreted by mesodermal cells.

### Axial organoids show sequential somite segmentation coordinated with traveling segmentation clock waves

Given that the morphology, composition, and signaling profiles of 120-hour-old organoids were consistent with those of mammalian embryos, we next measured the dynamics of somitogenesis. We tested whether the organoids showed anteriorly propagating segmentation clock waves in the PSM and a coordinated posteriorly propagating somite determination front. To do so, we performed time-lapse imaging of organoids built with the dual reporter HES7-Achilles/MESP2-mCherry iPSC line. After 72h of differentiation, when the first somite cells appeared at the anterior end, we transferred organoids to individual wells of a glass-bottom 96-well plate for imaging. Each organoid generated traveling waves of oscillating HES7 expression that were initiated at the posterior tip and moved anteriorly through the PSM (Fig. 3, A and B, Fig. S3B, Movie S1). When this wave reached the anterior end of the PSM, a new stripe of MESP2 expressing tissue was generated (Fig. S3A, Movie S2). Quantification of Achilles and mCherry signals showed that the MESP2+ region expanded in a step-like fashion, and each step in the MESP2 profile coincided with a peak of HES7 oscillations at the anterior end of the PSM, indicating that the timing of somite differentiation is coordinated with the segmentation clock wave in the axial organoids (Fig. 3, C and D, Fig. S3A). Characterization of the HES7 oscillations along the A-P axis revealed a global phase gradient, wherein oscillations at the anterior tip lagged the posterior tip by π/2 radians (n=14, Fig. 3E, fig. S3B). We also found that the period of oscillations was 4.5 hours and constant throughout the A-P axis (Fig. 3E). In mouse, chick, and zebrafish embryos, the NOTCH pathway has been implicated in driving intracellular gene expression oscillations and somite segmentation (Conlon et al., 1995; Dequeant et al., 2006; Oginuma et al., 2008), In line with *in vivo* studies (Ferjentsik et al., 2009; Niwa et al., 2011), inhibition of NOTCH signaling through DAPT (25μM) treatment resulted in downregulation of HES7 oscillation amplitude and the impairment of MESP2 progression, consistent with MESP2 being a NOTCH target gene (Fig. 3F, Fig. S3C, Movie S3). Thus, our *in vitro* organoid model recapitulates traveling segmentation clock waves and sequential somite segmentation as observed *in vivo*.

**Fig.3.**
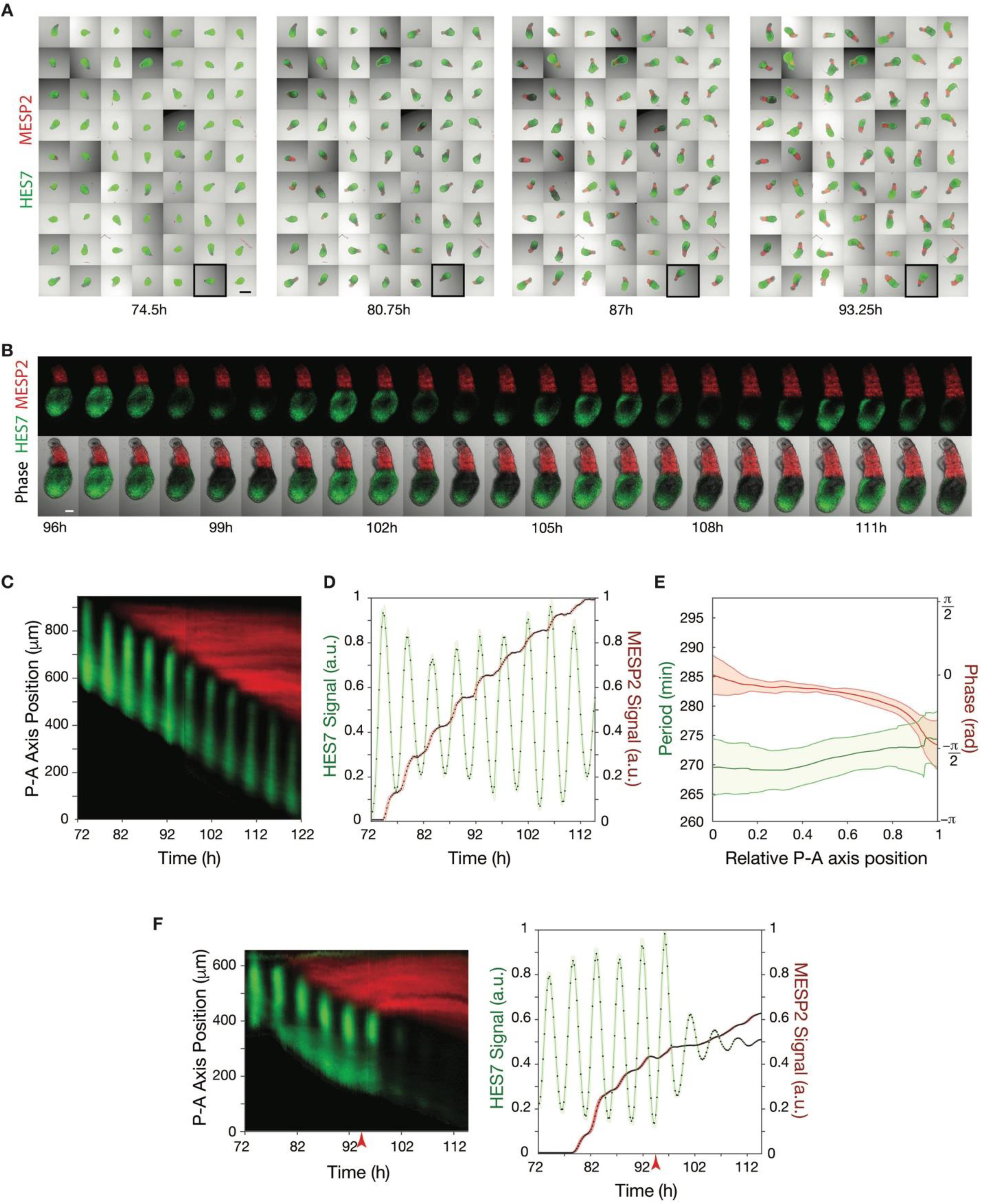
Dynamics of Somitogenesis and NOTCH gene expression waves in the organoids. (**A**) Stills from time-lapse imaging of organoids (at 74.5h, 80.75h, 87h and 93.25h after onset of differentiation) with HES7 (green) and MESP2 (red) expression reporters. All organoids show anteriorly moving oscillations of HES7 expression waves and posteriorly propagating MESP2+ somite determination front. Scale bars, 500µm. (**B**) Stills from time-lapse imaging of the organoid highlighted with a black box in (A) from 96h to 112.5h. Organoid shows anteriorly propagating HES7 (green) expression waves and posteriorly propagating MESP2 (red) somite determination front. Time interval between consecutive images is 45 minutes. Scale bar, 100µm. (**C**) Representative kymograph showing the dynamics of HES7 (green) and MESP2 (red) expression along the anteroposterior axis of organoids from 72 h to 122 h of differentiation. Data collected every 15 minutes. (**D**) HES7 (green) expression at the most anterior tip of the presomitic mesoderm and the length of the MESP2 (red) expressing region of a representative organoid over time. Data collected every 15 minutes. Thicker lines show the moving average with a window size of 3 timepoints and shaded areas show moving standard deviation with a window size of 3 timepoints. Black dots represent each data point. AU, arbitrary units. (**E**) Representative plot showing period (left axis) and phase (right axis) of the oscillations along the anteroposterior axis of presomitic mesoderm of organoids. Lines represent mean and shaded area represent standard deviation over n=14 biologically independent replicates. (**F**) Left: Representative kymograph showing the dynamics of HES7 (green) and MESP2 (red) expression along the anteroposterior axis of organoids from 72h to 114h of differentiation upon NOTCH inhibition using DAPT. Right: HES7 (green) expression at the most anterior tip of the presomitic mesoderm and the length of the MESP2 (red) expressing region of a representative organoid over time upon NOTCH inhibition. Data collected every 15 minutes for both plots. Thicker lines show the moving average with a window size of 3 timepoints and shaded areas show moving standard deviation with a window size of 3 timepoints. Black dots represent each data point. AU, arbitrary units. Red arrow indicates time of DAPT addition.

### FGF and WNT pathways have complementary roles in axial elongation, movement of the somite determination front and somite segmentation

In our organoids, WNT inhibition using the WNT secretion inhibitor IWP-2 (2μM) added during somitogenesis resulted in truncated organoids with truncated PSM (Fig. 4A and B, Fig. S4A). However, we did not observe an effect of WNT inhibition on the movement of the somite determination front, and we continued to see step-like segmentation with segment sizes similar to control organoids (Fig. 4C, Fig. S4A, Movie S4). Consistently, when ectopic WNT activation was uniformly stimulated by CHIR (3μM) addition at 96h of differentiation, organoids were elongated and developed a longer PSM compared to controls (Fig. 4D, Fig. S4, B and C). These organoids continued to show step like MESP2 progression without a significant change in somitic mesoderm length or defects in segmentation compared to control organoids (Fig. 4, D-F, Fig. S4, B and C, S4E). These results show that while the WNT pathway directly promotes axial elongation and generation of PSM, it has little effect on the progression of the determination front or somite segmentation.

**Fig.4.**
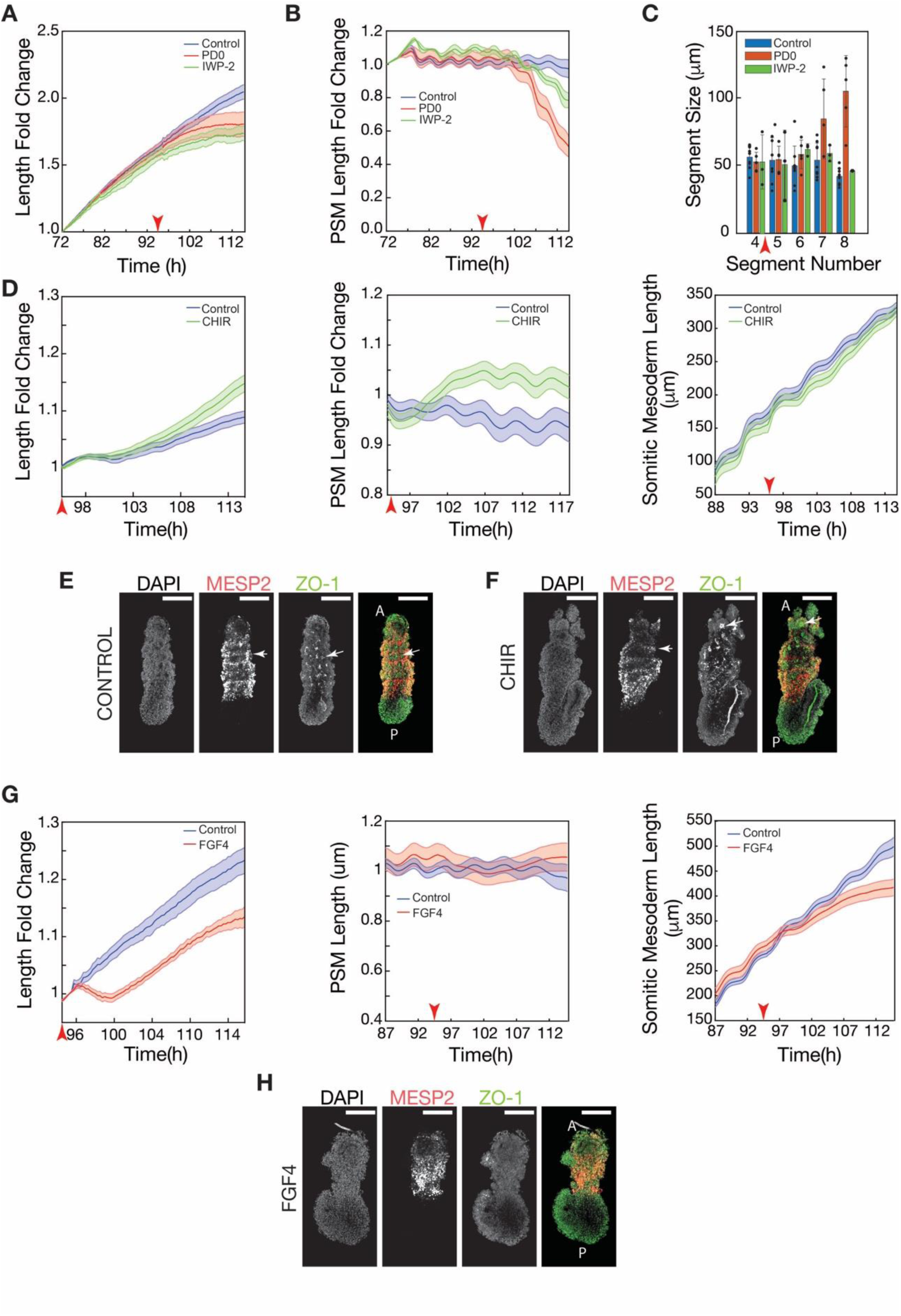
FGF drives somite differentiation front propagation and somite segmentation while WNT drives axial elongation. (**A** and **B**) Length fold change (A) and presomitic mesoderm length fold change (B) of organoids treated with PD0325901 (1µM, n=5, red), IWP-2 (2µM, n=4) and unperturbed control (n=13, blue) over time. Red arrow shows the timepoint of administration of PD0325901 and IWP-2 for the perturbed organoids. solid lines: mean, shaded areas: standard error. (**C**) Bar graph of somite segment sizes in organoids treated with PD0325901 (1µM, n=5), IWP-2 (2µM, n=4) and unperturbed control (n=9). Bars represent the mean; whiskers represent the standard deviation. Black dots represent individual organoids. (**D**) Length fold change (left), presomitic mesoderm length fold change (middle) and somitic mesoderm length (right) as a function of time in organoids treated CHIR (n=16, green) and unperturbed control (n=16, blue). Red arrow shows the timepoint of administration of CHIR for the perturbed organoids. solid lines: mean, shaded areas: standard error. (**E** and **F**) Confocal images of control (E) and CHIR treated (F) Typical images of organoids stained for DAPI, epithelial marker ZO-1 and somite marker MESP2, show MESP2 expression pattern resolved into an alternating pattern in each segment (one segment marked with white arrow) and epithelial segmented somites (ZO-1 puncta in one individual somite marked with white arrow). The CHIR treated organoid is substantially longer than control. Scale bar, 200µm **(G)** Length fold change (left), presomitic mesoderm length fold change (middle), somitic mesoderm length (right) as a function of time in organoids treated with FGF4 (n=9, red) and unperturbed control (n=13, blue). Red arrow shows the timepoint of administration of FGF4 for the perturbed organoids. solid lines: mean, shaded areas: standard error. **(H)** Typical image of an FGF4 treated organoid stained for DAPI, epithelial marker ZO-1 and somite marker MESP2. In contrast with control organoid in (E), FGF4 treated organoids do not show segmentation, and no ZO-1 puncta associated with epithelial somites. Scale bar, 200µm.

Conversely, experiments in mouse have indicated that WNT plays a role in defining the position of the determination front (Aulehla et al., 2008). These results were obtained in mice through β-catenin deletion or stabilization. Unlike experiments in organoids where the perturbation can be carefully timed, the effects of these mutations last throughout development, making it difficult to distinguish direct from indirect effects. Indeed, levels of FGF ligand and signaling activity were also significantly affected in these mutants, making it difficult to disentangle the effects of WNT perturbations from the downstream effects through FGF. Therefore, we next tested the role of FGF signaling in axial morphogenesis.

Consistent with studies in chick (Dubrulle et al., 2001) and zebrafish (Sawada et al., 2001) embryos, FGF inhibition in the organoids using FGF/ERK inhibitor PD0325901 (1μM) led to truncation, with both the total organoid length and PSM length being shorter than control (Fig 4, A and B, Fig. S4A). FGF inhibition further led to an accelerating somite differentiation front, and somite segments were about twice as large compared to control organoids (Fig. 4C, Fig. S4A Movie S5). To observe the effects of FGF activation during somitogenesis, we exposed organoids to FGF4 (100 ng/mL) ligand at 96 h of differentiation. Uniform FGF4 treatment did not change the size of the PSM but led to truncation of the organoid compared to control (Fig. 4G, Fig. S4D). FGF4-treated organoids had a decelerated determination front progression (Fig. 4G) and disrupted somite segmentation as seen by the lack of ZO-1 foci or segments in the perturbed organoids (Fig. 4H, Fig. S4E).

These results indicate that while both FGF and WNT pathways are important for the axial elongation of both the neural tube (Anand et al, 2022) and mesoderm, as well as the induction of PSM, the FGF pathway additionally plays a direct role in the definition of the somite determination front and in the segmentation of the somites.

### FGF4 gradients are required for the propagation of segmentation clock waves

How waves of gene expression travel anteriorly along the presomitic mesoderm remains unknown (Diaz-Cuadros and Pourquie, 2021). Furthermore, we do not know how the waves interact with the FGF and WNT signaling gradients. To determine if diffusible signaling gradients played a role in the propagation of segmentation clock waves, we wanted to place isolated colonies of PSM cells at a distance from each other to determine whether they could still communicate and mutually coordinate their segmentation clock oscillations through diffusive signals. This would imply that diffusive molecules were potentially important for the spatial coordination of the segmentation clock. To achieve this, we micropatterned pluripotent stem cell colonies in proximity to each other on a coverslip and constrained colony expansion and cell migration by passivating the coverslip surface (Fig. 5A, Fig. S5, A and B). The colonies could not touch each other, preventing any communication between colonies through juxtacrine signaling (such as through NOTCH or YAP). By treating cells with CHIR and LDN for 48 hours, we differentiated colonies to a presomitic mesoderm identity. At the end of 48 hours, we replaced differentiation media with basal media and recorded HES7 oscillations (Fig. 5A, Fig. S5A). If diffusible signals did not play a role in generating traveling waves, we expected not to see a phase gradient across the colonies and, concomitantly, no propagation of NOTCH activity waves sweeping coherently across the colonies. Contrarily, time-lapse imaging revealed that the HES7 oscillations were spatially coordinated between the microprinted colonies, and a phase gradient of HES7 oscillations was established, resulting in HES7 waves propagating from colonies at the edge to those in the center of the pattern (Fig. 5A-C, Figure S5A, Movie S6). This result led us to conclude that diffusible signals that caused inter-colony communication could be important for the propagation of segmentation clock waves.

**Fig.5.**
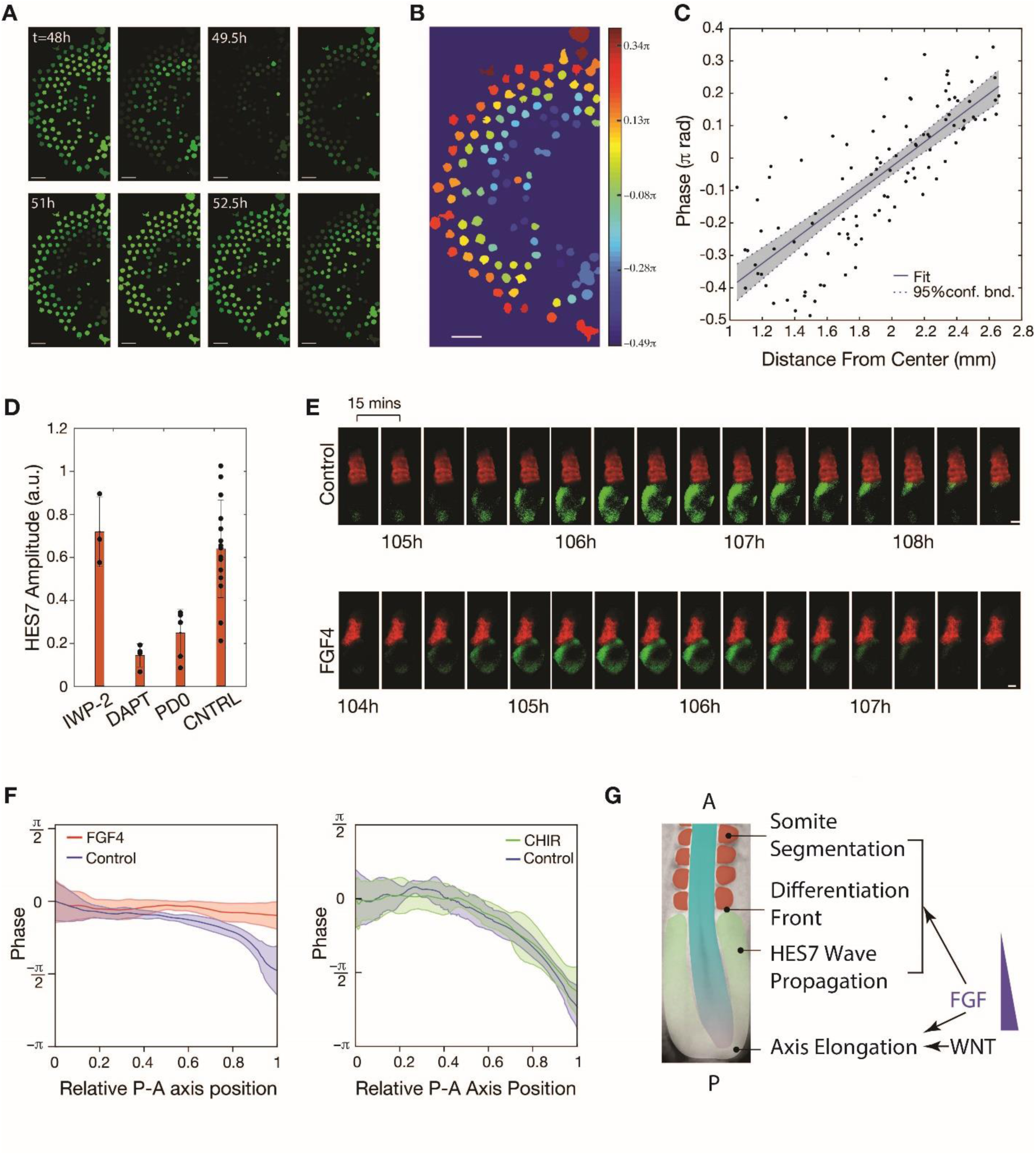
FGF gradient is required for HES7 traveling expression waves and somite segmentation. (**A**) Stills from time-lapse imaging PSM colonies on microcontact printed arrays with HES7 expression reporter. Detrended HES7 signal averaged over each colony is represented by green color intensity. Scale bar, 500 µm. (**B**) Colonies colored by their oscillation phase. Scale bar, 500 µm. (**C**) Plot showing oscillation phase of each colony and its distance from the center from the array. Black dots represent individual colonies. Blue line is obtained by linear regression of all data points. Dashed lines represent the 95% confidence bounds. Positive slope of the line indicates waves traveling from edge to inside of the array. (**D**) Bar graph of oscillations of HES7 amplitude of organoids treated with IWP-2 (2µM, n=3), DAPT (25µM, n=4), PD0325901 (1µM, n=5) and unperturbed control (n=15). Amplitude of the oscillations was calculated by normalizing the amplitude of 4^th^ peak after the treatment by the amplitude of 1^st^ peak after the treatment. Bars represent the mean; whiskers represent the standard deviation. Black dots represent individual organoids. (**E**) Stills from time-lapse imaging of organoids with HES7 (green) and MESP2 (red) expression reporters for (Top: Unperturbed Control, bottom: FGF4 addition). Time interval between images is 15 min. Scale bar, 100 µm. (**E**) Plots showing the phase profile of the organoids treated with FGF4 (left) and CHIR (right) compared to unperturbed control organoids. (see Methods). Lines represent mean and shaded area represent standard deviation over (Left: FGF4, n=9; Control, n=14, right: CHIR, n=10; Control, n=9) biologically independent replicates. (**J**) Proposed mechanism for A-P patterning of paraxial mesoderm wherein FGF controls wave propagation, somite segmentation and differentiation front propagation while WNT while FGF and WNT together control axial elongation.

We, therefore, investigated whether FGF and WNT signaling could modulate the propagation of the measured HES7 waves in organoids. We observed that FGF inhibition resulted in downregulation of the amplitude of HES7 oscillations, while WNT inhibition did not affect the oscillations (Fig. 5D), indicating that the FGF pathway is not only involved in defining the somite determination front and regulating somite segmentation, but also in the regulation of the segmentation clock. We next uniformly activated FGF and WNT pathways in organoids. Time-lapse imaging of organoids revealed that uniformly activating the WNT pathway by adding CHIR did not affect the phase gradient along the A-P axis compared to the control organoids (Fig. 5, E and F, Fig. S5, C and D, Movie S7) showing that the WNT pathway has no direct effects on the dynamics of the segmentation clock. Treating organoids with FGF4 ligand resulted in synchronization of the oscillations throughout the A-P axis and a loss of the phase gradient, in contrast to control organoids where segmentation clock waves propagated anteriorly through the PSM (Fig 5, E and F, Fig. S5E, Movie S8). Thus, we concluded that the FGF signaling gradient drives the propagation of segmentation clock waves during human somitogenesis.

## Discussion

In physical systems that break symmetry, by coupling the underlying degrees of freedom, one can substantially reduce the entropy of the broken symmetry state. Confirming the power of the approach developed in Anand *et al*., our study shows that by similarly coupling organoids, one can reduce the entropy of the broken symmetry state to obtain robust differentiation. The resulting axial organoids recapitulate key aspects of axial development including axial elongation, single-lumen neural tube formation, traveling segmentation clock waves, and sequential somite segmentation. These organoids allow us to image and deliver perturbations with temporal precision and at timescales corresponding to the clock oscillation times to extract mechanistic insight. Such an ability gives us an opportunity to study mammalian and in particular human axial development and associated diseases.

Our findings here and in Anand *et al*. show that WNT and FGF pathways together drive axial elongation. Unlike WNT, the FGF ligand gradient also defines the somite determination front and spatially modulates the activities of the NOTCH pathway and the segmentation clock to generate traveling gene expression waves (Fig. 5G). Loss of the FGF gradient through uniform activation of FGF signaling also led to somite polarity and segmentation defects. These results support the two-phase models (Jaeger and Goodwin, 2001; Oginuma et al., 2010) proposed to explain the somite anteroposterior polarity. In these models, the anteroposterior region of each somite is defined by the phase gradient of the segmentation clock in the presumptive somite region. Consistently, in our experiments, uniform FGF activation leads to the loss of both the phase gradient and anteroposterior polarity. Thus, by controlling both the anterior propagation of segmentation clock waves through the PSM and the posterior movement of the somite determination front, while independently driving axial elongation, the FGF pathway plays a central role in orchestrating the dynamics of axial patterning and morphogenesis in humans.

## SUPPLEMENTARY FIGURES AND MOVIE CAPTIONS

**Fig.S1.**
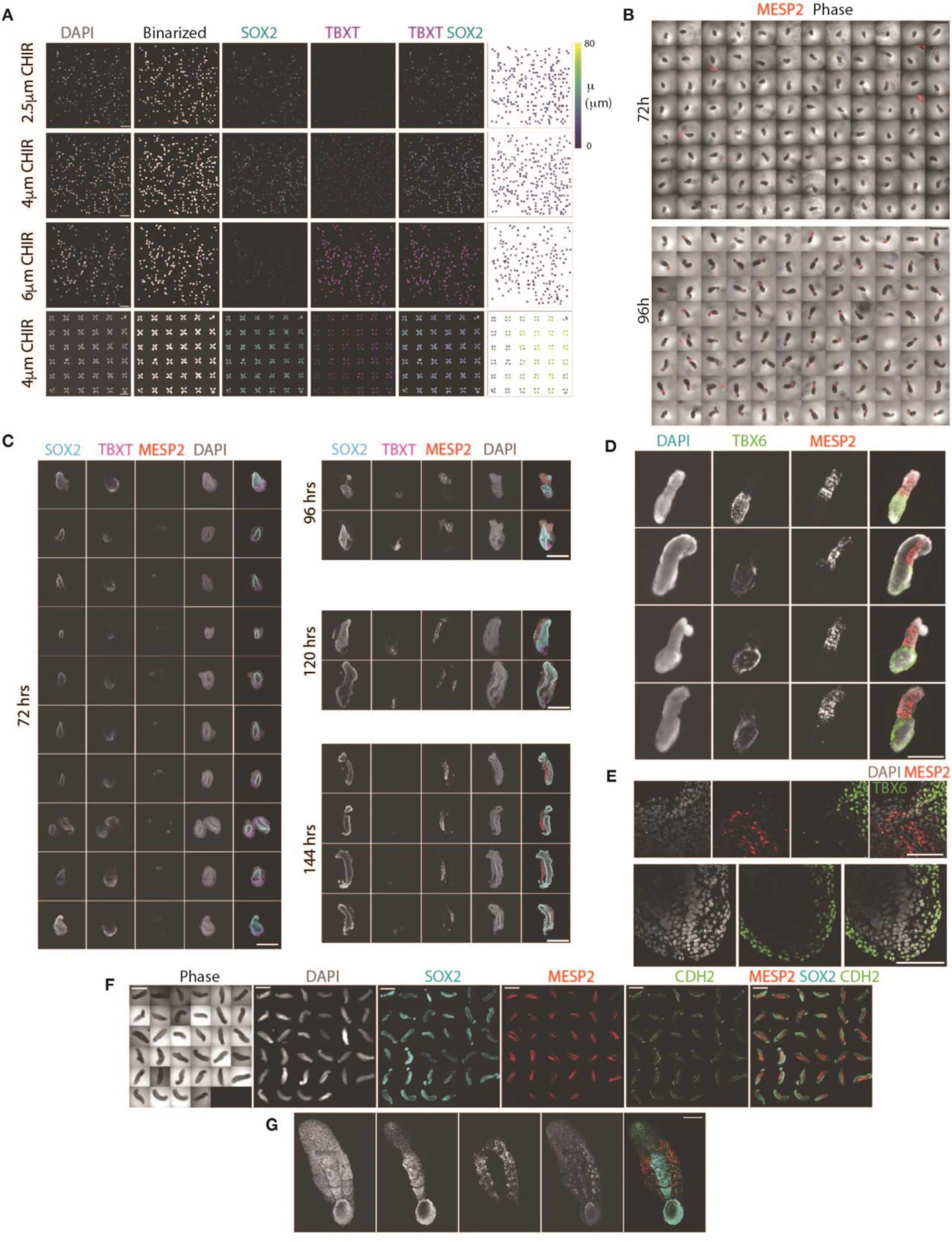
Elongating axial organoids generates neural tube with a single lumen flanked anteriorly by segmented somites and posteriorly by presomitic mesoderm. (**A**) Randomly positioned organoids (top three rows) and micropatterned organoids (bottom row) in groups of four on the vertices of a square on a coverslip, each consisting of a single epithelial layer of cells enclosing a single lumen, treated with BMP inhibitor LDN193189 (0.5µM), TGFβ inhibitor A83-10 (0.5µM) and WNT agonist CHIR99021 (top to bottom: 2.5µM, 4µM, 6µM, 4 µM) for 48 hours stained for DAPI, SOX2 and TBXT. Organoids were segmented based on DAPI signal. Rightmost column shows each organoid’s position on the respective row colored by their dipole moment. Scale bar, 1mm. (**B**) Phase contrast images overlayed with MESP2::mCherry signal in live organoids in a 96-well low adhesion plate at 72 h (top) and 96 h (bottom) of differentiation. Scale bars, 1 mm. (**C**) Confocal sections of representative organoids with MESP2::mCherry reporter on consecutive days of differentiation (72 h, 96 h, 120 h, 144 h) stained for SOX2, TBXT. SOX2 and TBXT co-expressing NMP’s reside at the posterior tip. Scale bars, 500 µm (**D**) Epifluorescence image of organoids with MESP2:mCherry reporter stained for paraxial mesoderm marker TBX6 at 120h of differentiation. Scale bars: 500µm. (**E**) Confocal sections of an organoid from with MESP2::mCherry reporter from (top) determination front and (bottom) posterior tip at 120 h of differentiation stained for TBX6. TBX6 and MESP2 expressing cells forms a clear boundary at the determination front. Scale bars, 100 µm. (**F**) Epifluorescence image of organoids with MESP2:mCherry reporter stained for neural marker SOX2 and N-Cadherin (CDH2) which is condensed at the apical side of epithelial cells. All organoids have a neural tube flanked by segmented epithelialized somites. Scale bars, 1 mm. (**G**) Maximum intensity projection from confocal sections of an organoid with MESP2::mCherry reporter stained for neural marker SOX2 and N-Cadherin (CDH2). Scale bars, 200µm.

**Fig.S2.**
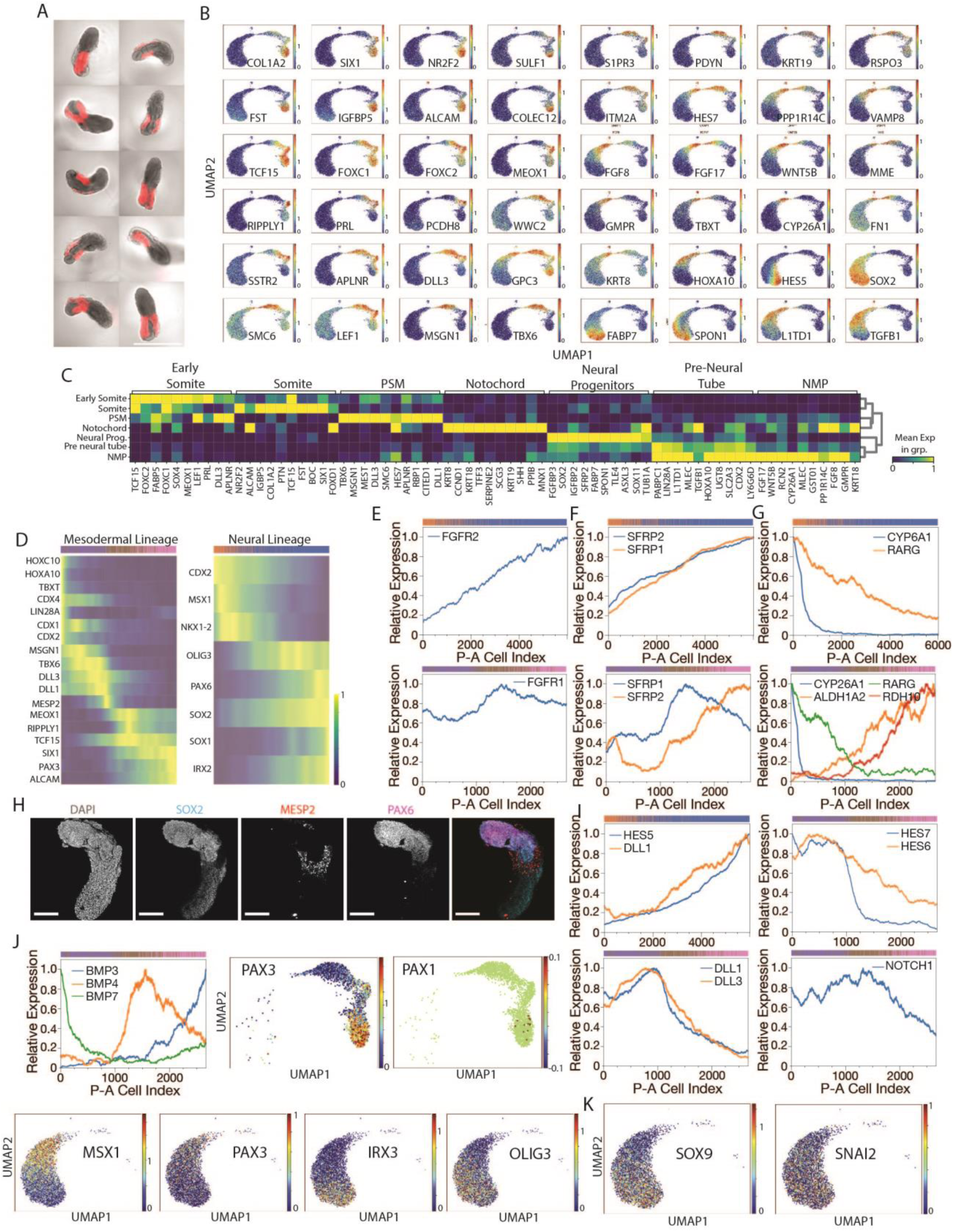
Anteroposterior organization of cell types and gene expression profiles inferred from single cell RNA-seq. (**A**) Epifluorescence image of organoids with MESP2:mCherry reporter at 120h of differentiation. We performed single-cell RNA sequencing of 11009 cells obtained from these organoids after dissociation and pooling. Scale bar: 1 mm. (**B**) UMAP (uniform manifold approximation and projection) plots showing the log-normalized gene expression values of the genes identified by SMD. (**C**) Heatmap of top 10 differentially expressed genes with highest fold change for each cell type compared to the other cell types. Genes are colored by their normalized mean expression in the respective cell type. Normalization is done by scaling log-normalized expression of each gene between 0-1. (**D**) Heatmap of top key genes (y axis) for mesodermal (left) cell clusters (presomitic mesoderm, early somite, and somite) and neural (right) cell clusters (pre-neural tube and neural progenitors) in cells (x axis) ordered according to their inferred anteroposterior positions. Genes are ordered based on the position of their peak expression on the inferred A-P axis. Color bars on the top of heatmaps represent the cluster identity of the individual cells (same color code as in Fig. 2A). (**E**) Normalized posterior-anterior gene expression profiles for FGFR2 in neural clusters (top), FGFR1 in mesodermal clusters (E, bottom) (**F**) Normalized posterior-anterior gene expression profiles for secreted WNT pathway inhibitors, SFRPs in neural (top) and mesodermal (bottom) clusters. SFRP’s show high expression in the anterior for both neural and mesodermal clusters. Color bars on the top of plots represent the cluster identity of the individual cells (same color code as in Fig. 2A). (**H**) Confocal sections of an organoid with MESP2::mCherry reporter on 120 h of differentiation stained for SOX2 and PAX6. PAX6 expression is upregulated anterior to the determination front. Scale bar, 200 µm. (**I**) Normalized posterior-anterior gene expression profiles for NOTCH target HES5 and NOTCH ligand DLL1 in neural clusters (top left); NOTCH targets (top right), NOTCH ligands (bottom left) and NOTCH receptor (bottom right) in mesodermal clusters. Color bars on the top of plots represent the cluster identity of the individual cells (same color code as in Fig. 2A). (**J**) (Top left) Normalized posterior-anterior gene expression profiles for BMP ligands expressed in the mesodermal clusters. Color bars on the top of plots represent the cluster identity of the individual cells (same color code as in Fig. 2A). UMAP plots showing the log-normalized gene expression values of the genes associated with dorsal (PAX3, MSX1, IRX3, OLIG3) and ventral (PAX1) cell identities. (**K**) UMAP plots showing the log-normalized gene expression values of neural crest markers SOX9 and SNAI2 on neural cell clusters.

**Fig.S3.**
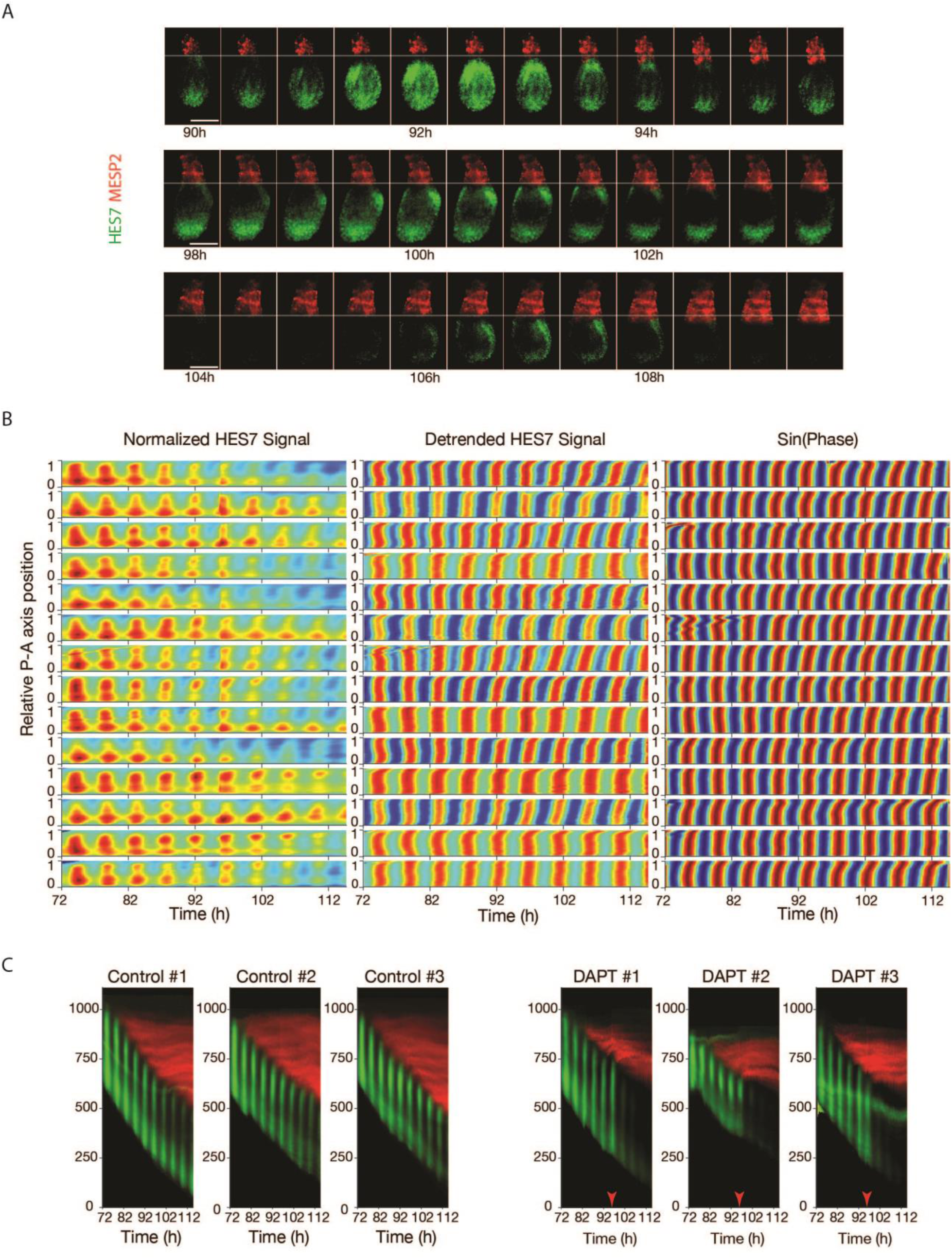
Dynamics of Somitogenesis and NOTCH gene expression waves in the organoids. (**A**) Stills from time-lapse imaging of three biologically independent organoids with HES7 (green) and MESP2 (red) expression reporters. Shaded line shows the position of the determination front at the first timepoint for each organoid. A new segment of MESP2 expression appears when each HES7 wave reaches to the determination front. Time interval between consecutive images is 30 minutes. Scale bars, 200 µm. (**B**) Kymographs of normalized HES7 signal (left), detrended HES7 signal (middle) and sine of the detected instantaneous phase of the oscillations for 14 organoids along the anteroposterior axis of organoids from 72 h to 122 h of differentiation. These kymographs were used to calculate the phase profile of Control organoids in Fig. 3E. Data collected every 15 minutes. For normalization, detrending of the signal and phase detection, see Methods. (**C**) Kymographs showing the dynamics of HES7 (green) and MESP2 (red) expression along the anteroposterior axis of organoids from 72 h to 114.75 h of differentiation for three control organoids (left) and three organoids treated with DAPT (25 µM) at 95h (right). Data collected every 15 minutes for all plots.

**Fig.S4.**
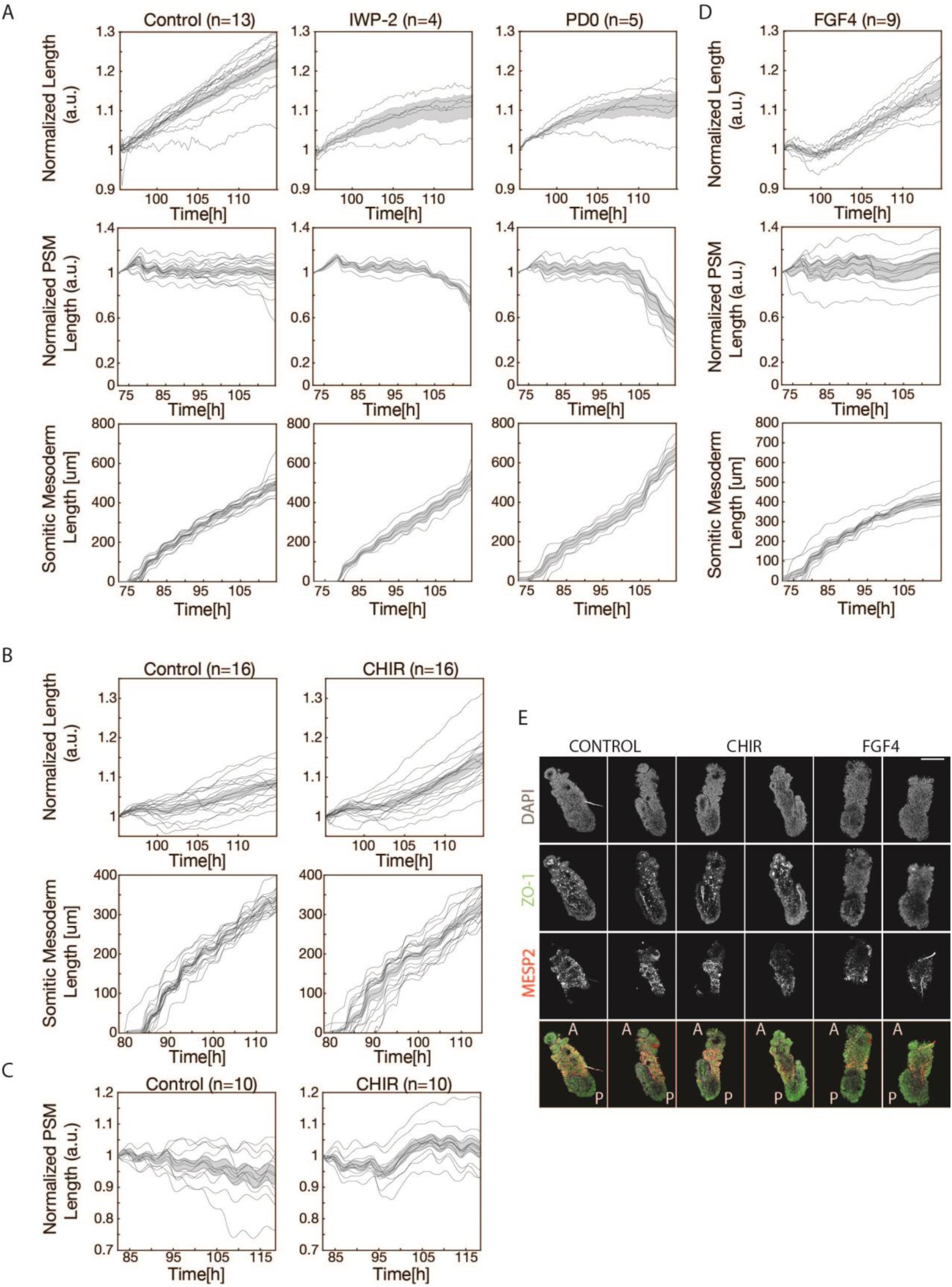
FGF drives somite differentiation front propagation and somite segmentation while WNT drives axial elongation. (**A**) Plots of length fold change (top row), presomitic mesoderm length fold change (middle row) and somitic mesoderm length (bottom row) of organoids treated with PD0325901 (1µM, n=5, right column), IWP-2 (2µM, n=4, middle column) and unperturbed control (n=13, left column) over time. Red arrow shows the timepoint of administration of PD0325901 and IWP-2 for the perturbed organoids. Solid lines represent individual organoids, shaded area: standard error around mean. (**B**) Plots of length fold change (top row), and somitic mesoderm length (bottom row) of organoids treated with CHIR (3µM, n=16, right column) unperturbed control (n=16, left column) over time. Red arrow shows the timepoint of administration of CHIR for the perturbed organoids. Solid lines represent individual organoids, shaded area: standard error around mean. (**C**) Plot showing presomitic mesoderm length fold change of organoids treated with CHIR (3µM, n=10, right) unperturbed control (n=10, left) over time. Red arrow shows the timepoint of administration of CHIR for the perturbed organoids. Solid lines represent individual organoids, shaded area: standard error around mean. (**D**) Plots of length fold change (top), presomitic mesoderm length fold change (middle) and somitic mesoderm length (bottom) of organoids treated with FGF4 (100 ng/mL, n=9) over time. Red arrow shows the timepoint of administration of FGF4 for the perturbed organoids. Solid lines represent individual organoids, shaded area: standard error around mean. (**E**) Confocal images of control (first two columns from left) CHIR treated (3^rd^ and 4^th^ column from left) and FGF4 treated (last two column from left) organoids at 120 h of differentiation stained for DAPI, epithelial marker ZO-1 and somite marker MESP2. Scale bar, 200µm.

**Fig.S5.**
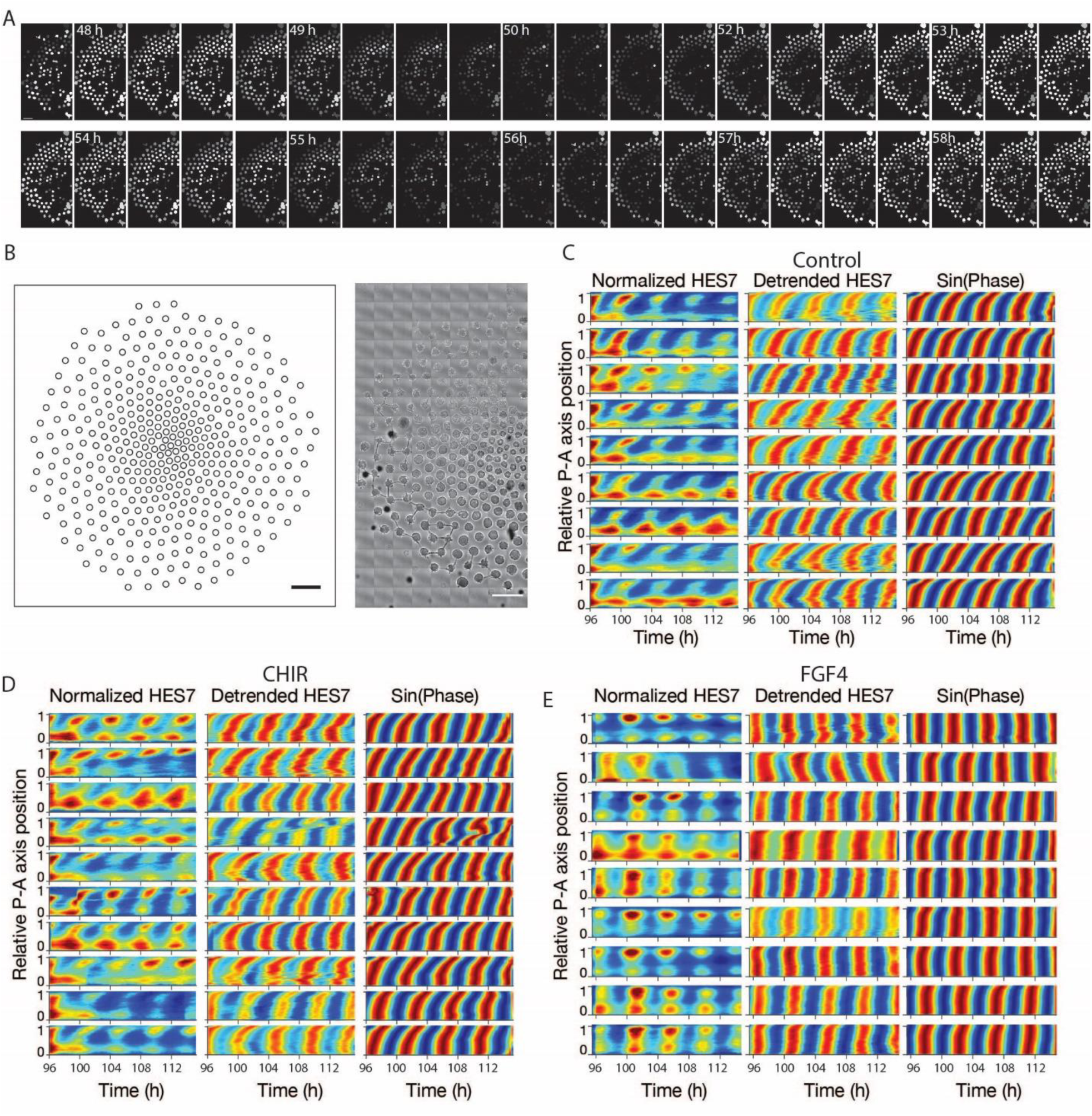
FGF gradient is required for HES7 traveling expression waves and somite segmentation. (**A**) Stills from time-lapse imaging PSM colonies on microcontact printed arrays with HES7 expression reporter. Detrended HES7 signal averaged over each colony is represented by green color intensity. Scale bar, 500 µm. (**B**) Left: Outline of the whole microcontact printed array. Each circle represents a colony. Scale bar, 1mm. Right: Phase contrast image of the microcontact printed colonies at 48 hours of differentiation. Scale bar, 500 µm. (**C-E**) Kymographs of normalized HES7 signal (left), detrended HES7 signal (middle) and sine of the detected instantaneous phase of the oscillations for 9 unperturbed organoids (C) 10 organoids treated with CHIR (D) and 9 organoids treated with FGF4 (E) along the anteroposterior axis of organoids. Kymographs in (C) and (D) were used to calculate the phase profile of Control and CHIR treated organoids in Fig. 5F, right, respectively. Kymographs in (E) were used to calculate the phase profile of FGF4 treated organoids in Fig. 5F, left. Data collected every 15 minutes.

### Supplementary Movie Captions

**Movie S1. Timelapse imaging of 63 organoids with HES7 (green) and MESP2 (red) expression reporters** 63 organoid in basal DGIP media supplemented with 6 %v/v Matrigel, between 72-90 hours of differentiation. HES7 and MESP2 expression was overlaid on phase contrast images. Scale bar, 1 mm. Time stamp shows the time passed after onset of differentiation.

**Movie S2. Timelapse imaging of a representative organoid with HES7 (green) and MESP2 (red) expression reporters** The organoid was imaged in basal DGIP media supplemented with 6 %v/v Matrigel, between 72-138 hours of differentiation. Time stamp shows the time passed after the differentiation was started. On the left, HES7 and MESP2 expression was overlaid on phase contrast images. Scale bar, 200 µm. Time stamp shows the time passed after the differentiation was started.

**Movie S3. Timelapse imaging of a representative organoid with HES7 (green) and MESP2 (red) expression reporters upon NOTCH inhibition**. The organoid treated with DAPT (25µM) at 95.5 hour of differentiation. On the left, HES7 and MESP2 expression was overlaid on phase contrast images. Scale bar, 200 µm. Time stamp shows the time passed after the differentiation was started.

**Movie S4. Timelapse imaging of a representative organoid with HES7 (green) and MESP2 (red) expression reporters upon WNT inhibition**. The organoid was treated with IWP-2 (2µM) at 95.5 hour of differentiation. On the left, HES7 and MESP2 expression was overlaid on phase contrast images. Scale bar, 200 µm. Time stamp shows the time passed after the differentiation was started.

**Movie S5. Timelapse imaging of a representative organoid with HES7 (green) and MESP2 (red) expression reporters upon FGF/ERK inhibition**. The organoid was treated with PD0325901 (1µM) at 95.5 hour of differentiation. On the left, HES7 and MESP2 expression was overlaid on phase contrast images. Scale bar, 200 µm. Time stamp shows the time passed after the differentiation was started.

**Movie S6. Timelapse imaging of HES7 signal from microcontact printed colonies**. Detrended and normalized HES7 expression was averaged over the colony. Scale bar, 1mm. Time stamp shows the time passed after the differentiation was started.

**Movie S7. Timelapse imaging of a representative organoid with HES7 (green) and MESP2 (red) expression reporters upon WNT activation**. The organoid was treated with CHIR (3µM) at 96 hour of differentiation. On the left, HES7 and MESP2 expression was overlaid on phase contrast images. Scale bar, 200 µm. Time stamp shows the time passed after the differentiation was started.

**Movie S8. Timelapse imaging of a representative organoid with HES7 (green) and MESP2 (red) expression reporters upon FGF activation**. The organoid was treated with FGF4 (100 ng/mL) at 95.5 hour of differentiation. On the left, HES7 and MESP2 expression was overlaid on phase contrast images. Scale bar, 200 µm. Time stamp shows the time passed after the differentiation was started.

## Acknowledgments

We thank Nicole El Ali, Claire Reardon, and the Bauer Core at Harvard University for their work on the RNA sequencing used in this manuscript as well as for their expertise and assistance with flow cytometry and FACS. We thank the Douglas Richardson and the Harvard Center for Biological Imaging for help and advice. We thank the Weitz lab and Perry Ellis for advice and help with microfabrication. We thank members of the Ramanathan Lab and in particular Giridhar Anand, Deniz Cihat Aksel, Theresa Weis, William Weiter and Mustafa Basaran for their help with experiments and advice. We thank Margarete Diaz-Cuadros and the Pourquie lab for sharing the HES7:Achilles/MESP2:mCherry iPSC line used for imaging. We thank Olivier Pourquie, Richard Losick and Jessica Whited for advice and valuable discussions. This work was supported in part by NIH R01GM131105, R01MH123948 and by start-up funds from Harvard University.

## Ethical compliance

We used hESCs and hiPSCs in accordance with approvals by Harvard University IRB (protocol #IRB18-0665) and Harvard University ESCRO (protocol E00065).

## Data availability

The single cell RNA-seq dataset generated in this study will be made available in the NCBI GEO repository and the NCBI SRA by the time of publication, or in advance of that date upon reasonable request. All other data are available upon reasonable request.

## Code availability

All custom code used in this study is available from the authors upon reasonable request.

## Supplementary Materials

Figs. S1 to S5

Movies S1 to S8

